## Supplemental Figures and Legends for "Learning enhances behaviorally relevant representations in apical dendrites"

**Supplementary Movie Legends**

**Supplementary Movie 1. Example two-photon microscopy movie during behavioral session.**

Playback speed is in real time. “CS+” and “CS-” denote times of stimulus onset. 433 x 433 μm field of view.

**Supplementary Movie 2. Example SCAPE microscopy movie during behavioral session.** *Top*, maximum intensity projection (MIP) across the dorsal-ventral dimension showing horizontal extent of dendritic activity. *Bottom*, MIP across the medial-lateral dimension showing vertical extent of dendritic activity. Playback speed is in real time. “CS+” and “CS-” denote times of stimulus onset. 300 × 1050 × 234 μm field of view.

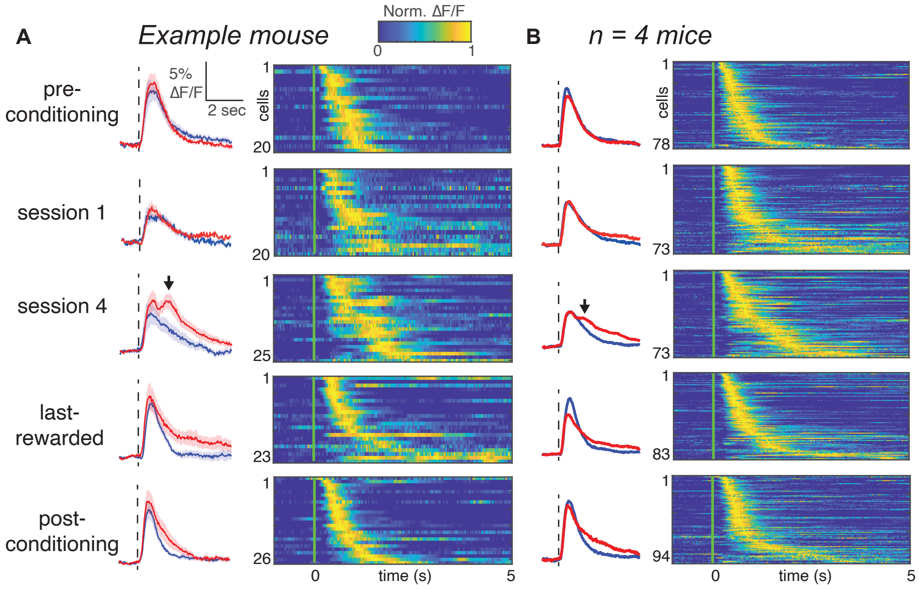


**Supplementary Fig. 1 I CS+ trials evoke a second, long-latency peak during early learning, but not late learning. a,** Left: Population average of stimulus-responsive tufts aligned to CS+ (red) or CS- (blue) trials from an example mouse. Right: Normalized ΔF/F of individual tufts during CS+ trials. **b,** Same as in **a**, combining data across four mice whose imaging regions were mapped with intrinsic imaging.

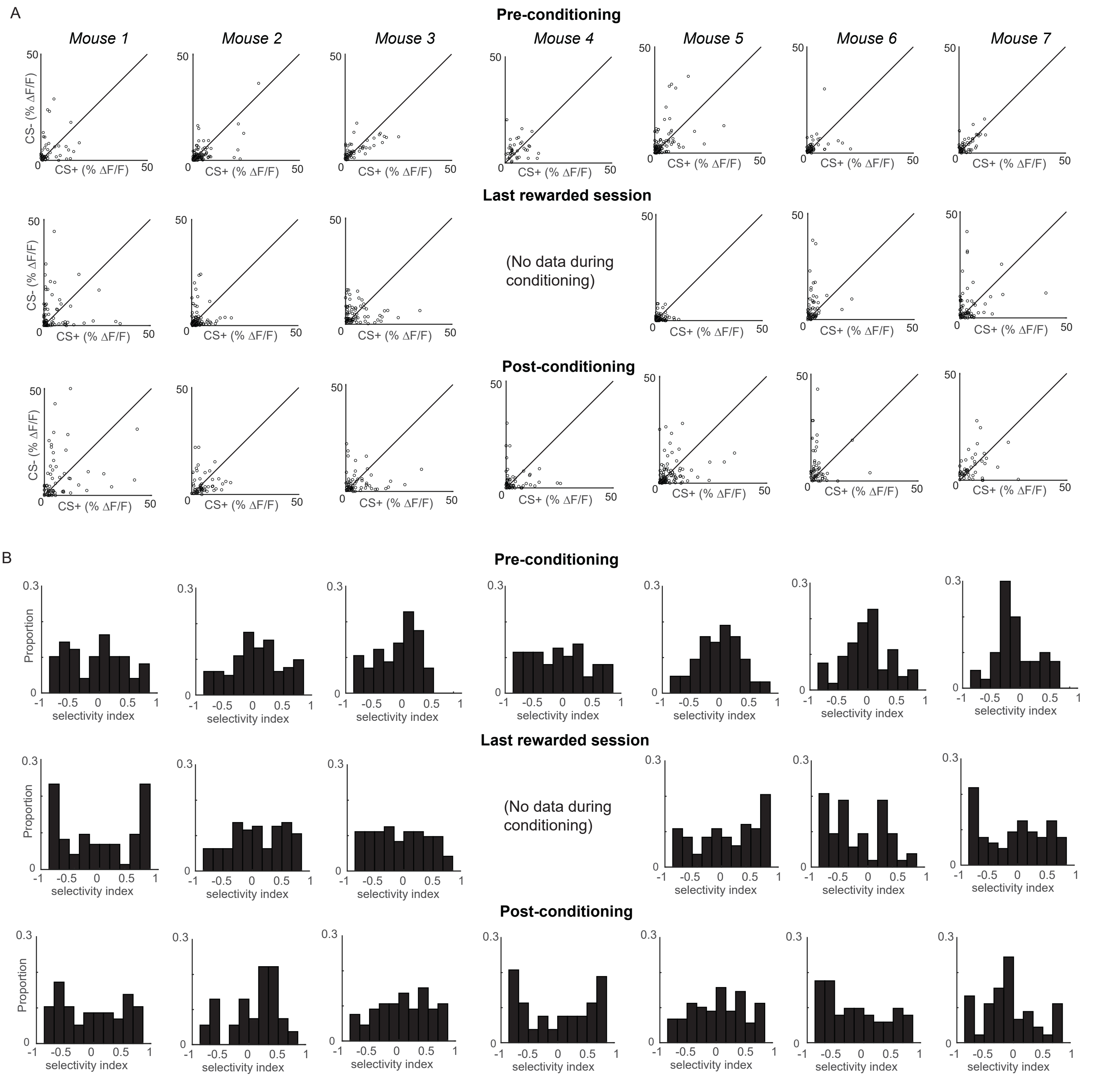


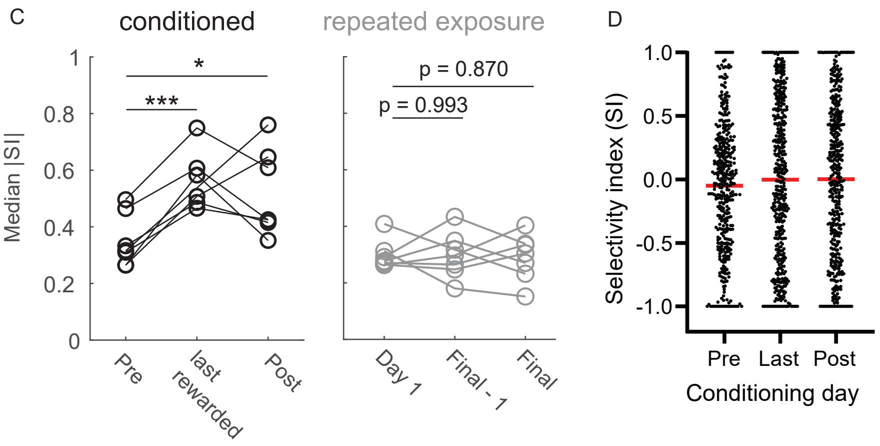


Supplementary Fig. 2 | Selectivity was enhanced in individual animals that received rewards. a, Tuft responses to CS+ and CS- in each conditioned animal. b, Selectivity indices for all tufts in ieach conditioned animal. c, Median SI magnitude for each animal across three sessions for conditioned (left) and repeated exposure groups (right). d, Scatter plot of the SI of all tufts on the Pre, Last, and Post conditioning days. Red lines indicate means. * $p < 0.05$, *** $p < 10^{-3}$

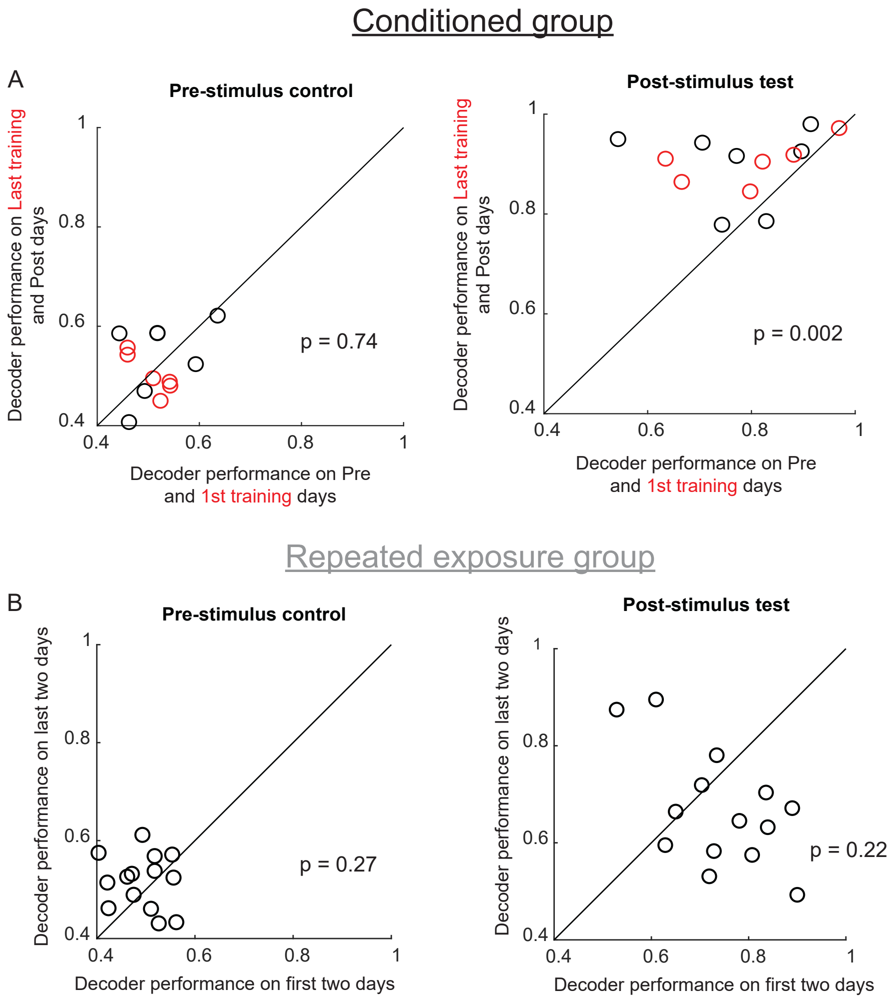


Supplementary Fig. 3 | Decoder performance improves after conditioning, but not repeated exposure. a, Decoder performance for each animal on the Pre and 1st training days was compared with performance on the Post and Last training days, respectively (thus two datapoints are shown for each animal. Red points indicate rewarded training sessions). Left: Decoder performance was low both before and after learning when trained on the pre-stimulus epoch. Right: Decoder performance improves after conditioning when trained on the post-stimulus epoch. b, Same analysis as A, but for the repeated exposure group. Decoder performance does not improve over time (right).

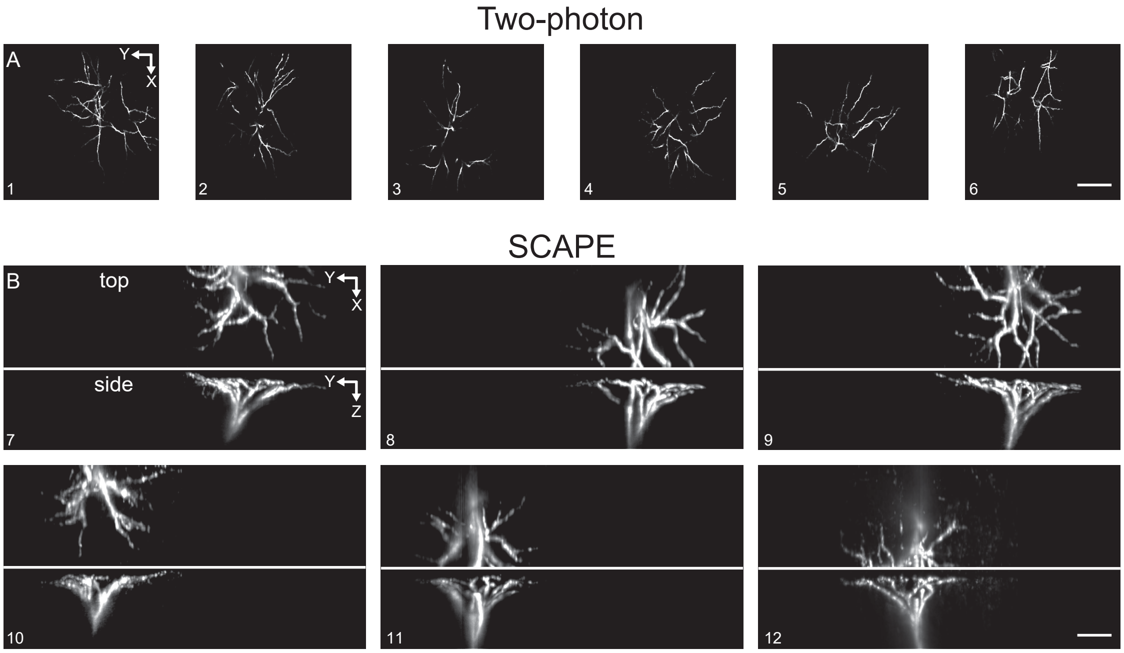


Supplementary Fig. 4 | Segmented tufts from two-photon and SCAPE microscopy. a,b, 12 example tufts extracted from either two-photon (a) or SCAPE microscopy (b). Tufts segmented from SCAPE microscopy are shown as maximum intensity projections from the top and side. Scale bars: 100μm.

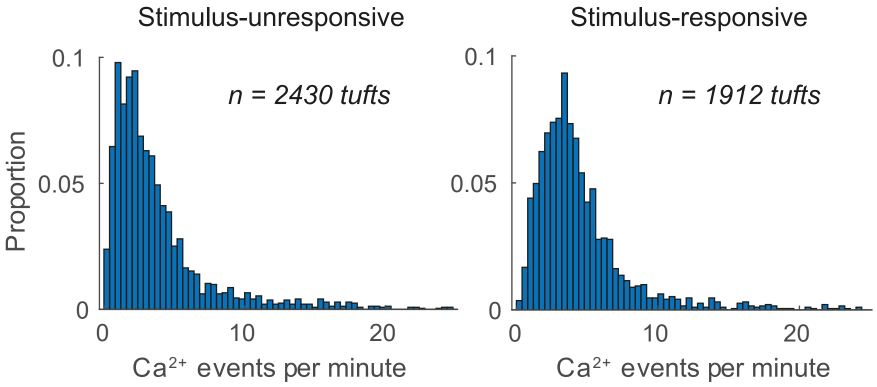


Supplementary Fig. 5 | Calcium event rate of tufts that were either unresponsive or responsive to air puff stimuli. The number of calcium events per minute was quantified for all tufts during each conditioning session. Data from each group was pooled across all sessions.

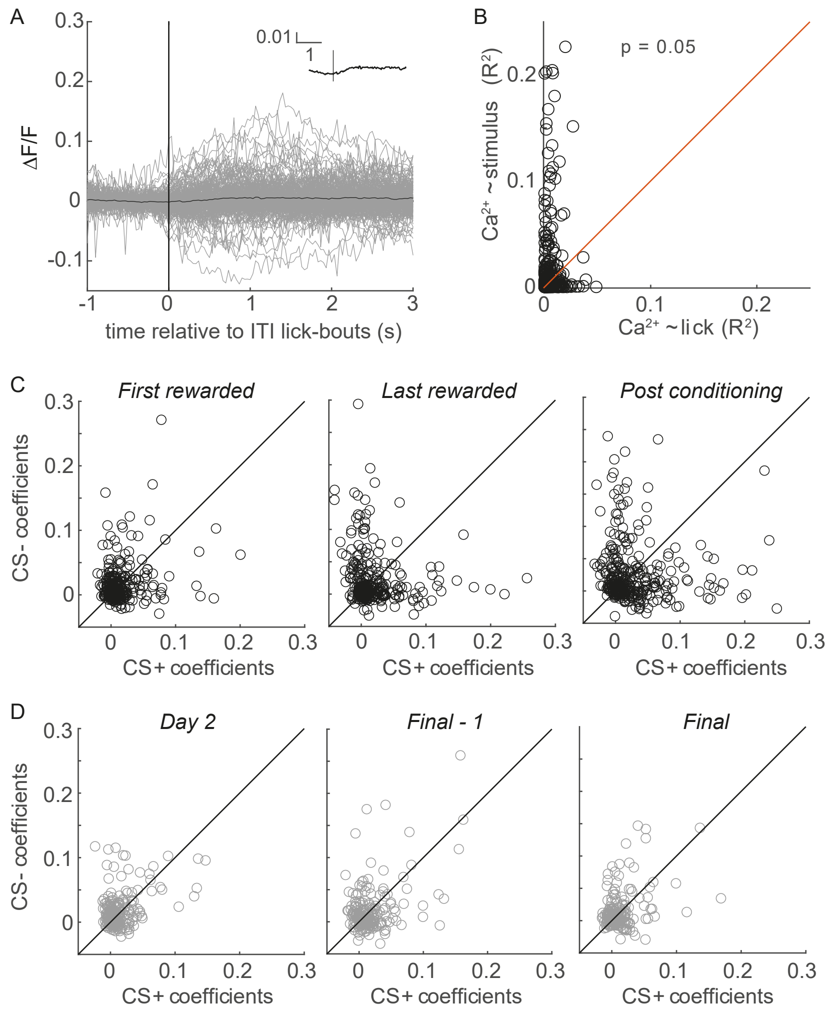


Supplementary Fig. 6 | Licking cannot account for changes in selectivity during learning. a, ITI lick-bout-triggered averages of 232 tufts on the 5th conditioning day, when ITI licks were still common (grey traces - individual tufts, black inset - population average), exhibit little or no lick-related calcium influx. b, R2 values for linear models predicting calcium from stimuli (y axis) are consistently greater than those predicting calcium from licking (x axis). Each circle represents one tuft out of 442 tufts on last-rewarded sessions. c, Coefficients from a multivariate regression analysis with calcium as the response variable and the CS+, CS-, whisking, and licking as the predictors. CS+ and CS- coefficients are therefore disentangled from correlations with whisking and licking. Conditioning biases individual tufts (circles) to have larger CS+ or CS- coefficients. n = 304, 324, and 322 tufts for First rewarded, Last rewarded, and Post conditioning, respectively. d, Similar analysis to C but for repeated exposure group, with calcium as the response variable and the CS+, CS-, and whisking as the predictors. n = 223, 208, and 218 tufts for Day 2, Final - 1, and Final session, respectively.
